# Snapshot of the evolution and mutation patterns of SARS-CoV-2

**DOI:** 10.1101/2020.07.04.187435

**Authors:** Jin Zhao, Jiumeng Sun, Wan-Ting He, Xiang Ji, Qi Gao, Xiaofeng Zhai, Marc A. Suchard, Samuel L. Hong, Guy Baele, Shuo Su, Jiyong Zhou, Michael Veit

## Abstract

The COVID-19 pandemic is the most important public health threat in recent history. Here we study how its causal agent, SARS-CoV-2, has diversified genetically since its first emergence in December 2019. We have created a pipeline combining both phylogenetic and structural analysis to identify possible human-adaptation related mutations in a data set consisting of 4,894 SARS-CoV-2 complete genome sequences. Although the phylogenetic diversity of SARS-CoV-2 is low, the whole genome phylogenetic tree can be divided into five clusters/clades based on the tree topology and clustering of specific mutations, but its branches exhibit low genetic distance and bootstrap support values. We also identified 11 residues that are high-frequency substitutions, with four of them currently showing some signal for potential positive selection. These fast-evolving sites are in the non-structural proteins nsp2, nsp5 (3CL-protease), nsp6, nsp12 (polymerase) and nsp13 (helicase), in accessory proteins (ORF3a, ORF8) and in the structural proteins N and S. Temporal and spatial analysis of these potentially adaptive mutations revealed that the incidence of some of these sites was declining after having reached an (often local) peak, whereas the frequency of other sites is continually increasing and now exhibit a worldwide distribution. Structural analysis revealed that the mutations are located on the surface of the proteins that modulate biochemical properties. We speculate that this improves binding to cellular proteins and hence represents fine-tuning of adaptation to human cells. Our study has implications for the design of biochemical and clinical experiments to assess whether important properties of SARS-CoV-2 have changed during the epidemic.

## Introduction

Coronavirus disease 2019 (COVID-19), caused by a *Betacoronavirus* named Severe Acute Respiratory Syndrome Coronavirus 2 (SARS-CoV-2), was initially reported in China in late December 2019^1,2^. As of the 20^th^ of Jun 2020, 8,525,042 infections and 456,973 deaths had been reported worldwide (data from WHO Situation Report). SARS-CoV-2 is a positive stranded enveloped RNA virus with a genome size of approximately 30 kb^1,3^. The virus genome comprises 14 open reading frames (ORFs) encoding 27 proteins. Virus particles are composed of four structural proteins: the Spike (S) protein, which is required for virus entry, the main determinant of host tropism and the target for neutralizing antibodies, the envelope (E) protein (an ion channel) and the matrix (M) protein are inserted into the viral envelope, whereas the nucleocapsid (N) protein is internal and enwraps the viral genome^4^. Some of the 8 accessory proteins (ORFs 3a, 3b, p6, 7a, 7b, 8b, 9b, 14) might also be virus components, but their functions are largely unexplored^3^. The 16 non-structural proteins (nsp1 to nsp16) are generated by proteolytic cleavage from two polyprotein precursors, pp1a and pp1ab, which are encoded by ORF1a and ORF1ab, respectively. ORF1ab accounts for more than 2/3 of the whole genome and is generated by ribosomal frame shifting during translation. The enzymes required for polyprotein processing are encoded by nsp3, a large multifunctional protein having a papain-like protease domain, and by the main protease (nsp5, 3C-like protease), which cleaves pp1a and pp1ab at eleven sites^5,6^. Most nsp proteins assemble into a membrane-bound replication and transcription complex. Anchoring of the complex to double-membrane vesicles that are derived from the endoplasmic reticulum membrane is achieved by the transmembrane proteins nsp4 and nsp6. The main enzymes for RNA processing are the RNA-dependent RNA polymerase (RdRp, nsp12) and the helicase (nsp13), but they are assisted by nsp7 to nsp16. Nsp1 and nsp2 are thought to modulate host cell responses^7,8^. Thus, most of the proteins encoded by ORF1a and ORF1ab are essential for virus replication and - at least for *Deltacoronaviruses* - for adaption of the virus to a new host^9^.

RNA viruses are known to be rapidly evolving, which could lead to the accumulation of amino acid mutations that might affect the transmissibility of the virus, its cell tropism and pathogenicity. Fortunately, until now, the observed diversity among pandemic SARS-CoV-2 sequences is low, but its rapid global spread may provide the possibility of positive natural selection. The lack of pre-existing immunity in the population and the high transmissibility of SARS-CoV-2 render its spontaneous disappearance unlikely^10,11^. Furthermore, it is not known whether SARS-CoV-2 is already fully adapted for efficient growth in human cells after its host-jump from bats or from a putative intermediate host^2,12^. This is usually associated with adaptive mutations as the analysis of genomes from early, middle and late cases during the SARS-CoV-1 epidemic revealed^13,14^. For avian H5N1 Influenza viruses it was shown that transmissibility of the virus is mainly associated with mutations in one polymerase protein and in the hemagglutinin (HA); the latter increased the stability of the molecule and its receptor binding affinity^15,16^. HA is also the target of antibodies, but during the flu season mutations slowly accumulate in HA and antibody escape mutants are generated that become the predominant strain^17^. Therefore, insights from the evolution of SARS-COV-2 will provide a better understanding of the epidemic and offer important implications for the development of vaccines.

So far, some studies have reported that SARS-COV-2 has evolved into new genotypes/subtypes, but those analyses of the phylogenetic tree were based on only a few hundred sequences^18-20^. Here, we developed an early warning pipeline to study SARS-CoV-2 evolution within the context of the pandemic. We used 4,894 SARS-CoV-2 sequences available from GISAID and NCBI GenBank (data release: April 11, 2020) to perform phylogenetic analysis based on whole genomes. In particular, we identified and characterized the geographic and temporal patterns of high-frequency mutations. Some of the mutations exhibit a dynamically changing pattern in their frequencies at different geographic regions which might indicate a potential signal of positive selection in the genes encoding ORF1ab and the structural proteins.

## Materials and Methods

### Sequence collection

A total of 6,193 complete genomic sequences of SARS-CoV-2 covering a time range from December 24, 2019 (the first isolate) to April 11, 2020 were downloaded from GISAID (https://www.gisaid.org/), including detailed sequence information (accession ID, virus name, location and collection date). Sequences with more than 300 unidentified bases were considered as low quality and were filtered out as well as sequences from cell passage isolates. The remaining 4,894 high-quality sequences which are at least 28,000bp in length, exhibit high coverage and were subsequently used for our analyses.

### Sequence Alignment

The 4,894 complete genome sequences of SARS-CoV-2 were aligned with MAFFT v7^21^ using the G-INS-i strategy and manually revised by using MEGA 7.0^22^. After aligning these genomes, we extracted the ORF1a (13218bp), ORF1ab (8072bp), S (3822bp), ORF3a (828bp), E (228bp), M (669bp), ORF6 (186bp), ORF7a (366bp), ORF7b (132bp), ORF8 (366bp), N (1260bp) and ORF10 (117bp) from each genome based on the reference strain (Accession NC_045512.2), and translated them into the corresponding amino acid sequences. For the next analysis, the twelve amino acid sequence sets were aligned by MEGA 7.0 using MUSCLE (Codons).

### Phylogenetic analysis of genome sequences, ORF1ab and S gene of SARS-CoV-2

The 4,894 complete genome, ORF1ab and S genes were used to reconstruct the maximum-likelihood (ML) phylogenetic tree using IQ-TREE v2.0.2^23^. The most suitable nucleotide substitution model was selected by the automatic model selection functionality in IQ-TREE^24^. In order to perform a more thorough search through tree space, we used both default IQ-TREE settings as well as additional parameters to intensify the search algorithm (allnni, ntop=100 and nbest=20). Multiple independent IQ-TREE analyses with these settings were performed and the tree with the highest (log) likelihood among these runs was retained. UltraFast bootstrap analyses were performed using 1,000 bootstrap trees. Trees were displayed by iTOL (https://itol.embl.de/) and colored by continent and country of each sequence^25^. The clusters were manually divided according to the topology of the whole genome phylogenetic tree, and then the genetic distance between clusters was calculated by MEGA 7.0 using the function “Compute Between Group Mean Distance” with the following parameters: Variance Estimation Method is none, the Substitutions Type is nucleotide, the Model/Method is p-distance and other parameters were default.

### Evolutionary analysis by determination of the ratio of non-synonymous versus synonymous nucleotide substitutions

The HyPhy software package was used to estimate the ratio of non-synonymous substitutions versus synonymous substitutions (dN/dS) and to identify the sites that are subjected to potential positive selection^26^. Phylogenetic trees of ORF1ab and S gene were used as reference trees in selection analysis, because ORF1ab has the most high-frequency mutation sites, and plays, besides S, an important role for virus replication. The following methods were used for selection analysis: SLAC (Single Likelihood Ancestry Counting), FEL (Fixed Impact Probability), MEME (Mixed Evolutionary Model of Evolution) and FUBAR (Fast Unconstrained Bayesian Approximation)^27-30^. We set stricter standards than usual for statistical significance, the *p* values of SLAC, FEL and MEME ≤ 0.05 and FUBAR posterior probability ≥ 0.95.

### Structural analysis

The software PyMol (https://pymol.org/2/) was used to create the figures from the pdb files. The integrated measuring wizard was used to determine the distance between two atoms. The electrostatic surface potential was calculated with the APBS plugin. Visualization of hydrophobicity and charge on protein surfaces was done with the YRB script^31^. Exchange of certain amino acids was performed with the integrated mutagenesis tool. Among the different possible rotamers of the mutated amino acid side chain the one was chosen that exhibits no clashes with neighboring amino acids. Prediction of phosphorylation sites was done with the NetPhos 3.1 tool (http://www.cbs.dtu.dk/services/NetPhos/).

## Result

### 1. Evolution of SARS-COV-2 and high-frequency amino acid mutations in SARS-COV-2 structural and non-structural proteins

We obtained 4,894 whole genome sequences from 60 countries and 6 continents on which we performed phylogenetic analysis (Fig. 1 and Fig. S1). The whole genome phylogenetic tree can be divided into five clusters/clades based on the tree topology and clustering of mutations. In these five clusters, the high-frequency mutation sites (with mutations greater than 1% in our data set of 4894 sequences) in each cluster are almost entirely different(Fig. S1). Each cluster also has a different geographical distribution: cluster I mainly consists of sequences from Asia and Europe, cluster II is mainly composed of sequences from North America and Asia. In cluster III, sequences from North America and Europe are alternately distributed, whereas clusters IV and V are dominated by European sequences. We found that the sequences from China gathered in clusters I and II, and the sequences from the United States clustered in clusters II and III. Sequences from European countries, such as the United Kingdom exist in each cluster, but are more prominently found in clusters I, IV and V. This indicates that there are differences in epidemic isolates of different regions. Of the five clusters, clusters IV and V have the smallest genetic distance to each other in the range of 10^−5^ substitutions per site. Cluster I contains the early Chinese sequences and all sequences from 2019, and exhibits the largest genetic distance to cluster II, which, however, is also small, in the range of 10^−4^ substitutions per site. It is worth noting that although the clustering of the whole genome phylogenetic tree is relatively clear visually and each cluster has its own characteristic mutation sites, the between group mean p-distance of five clusters is too small (Table S1) and its bootstrap value is too low to support genotyping, therefore, we did not perform more meticulous genotyping (Fig. S1).

**Figure 1:**
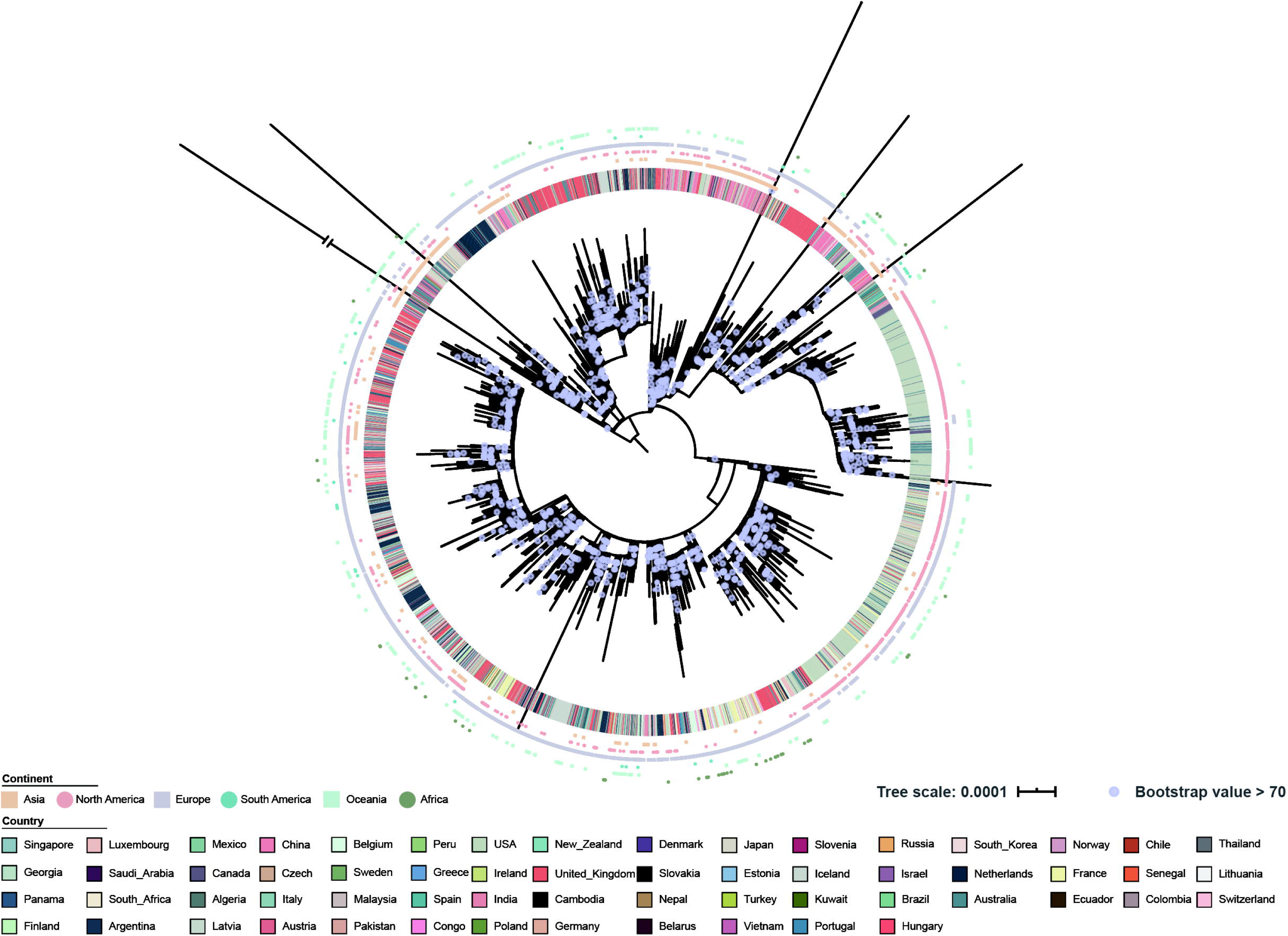
Geographical distribution and whole genome phylogenetic trees. Whole genome phylogenetic tree. The outer circle represents the origin of a sequence from a certain continent, and the inner circle from a certain country as specified in the inset.

The ORF1ab and whole genome phylogenetic trees have a lot in common. Comparing the structures of the trees revealed that the ORF1ab phylogenetic tree (Fig. 2) is visually similar to the whole genome phylogenetic tree, but the S protein phylogenetic tree is quite different and exhibits very low diversity and also low bootstrap support on internal nodes (Fig. S2). In terms of national distribution, the whole genome and ORF1ab phylogenetic tree show country aggregation, whereas the S protein phylogenetic tree is relatively scattered. This may be due to the high homogeneity of the S protein sequences, resulting in poor diversity. Like the whole genome tree, the ORF1ab and S gene phylogenetic trees do not support genotyping because of the low bootstrap values. It needs to be explained that the S protein phylogenetic tree exhibits low bootstrap support and low diversity, and has no significant features except for the Asp614Gly mutation site that has been reported many times. This shows that there is limited genetic variation in the currently sampled viruses, but more recent ones are showing more divergence in location as expected for fast evolving RNA viruses.

**Figure 2:**
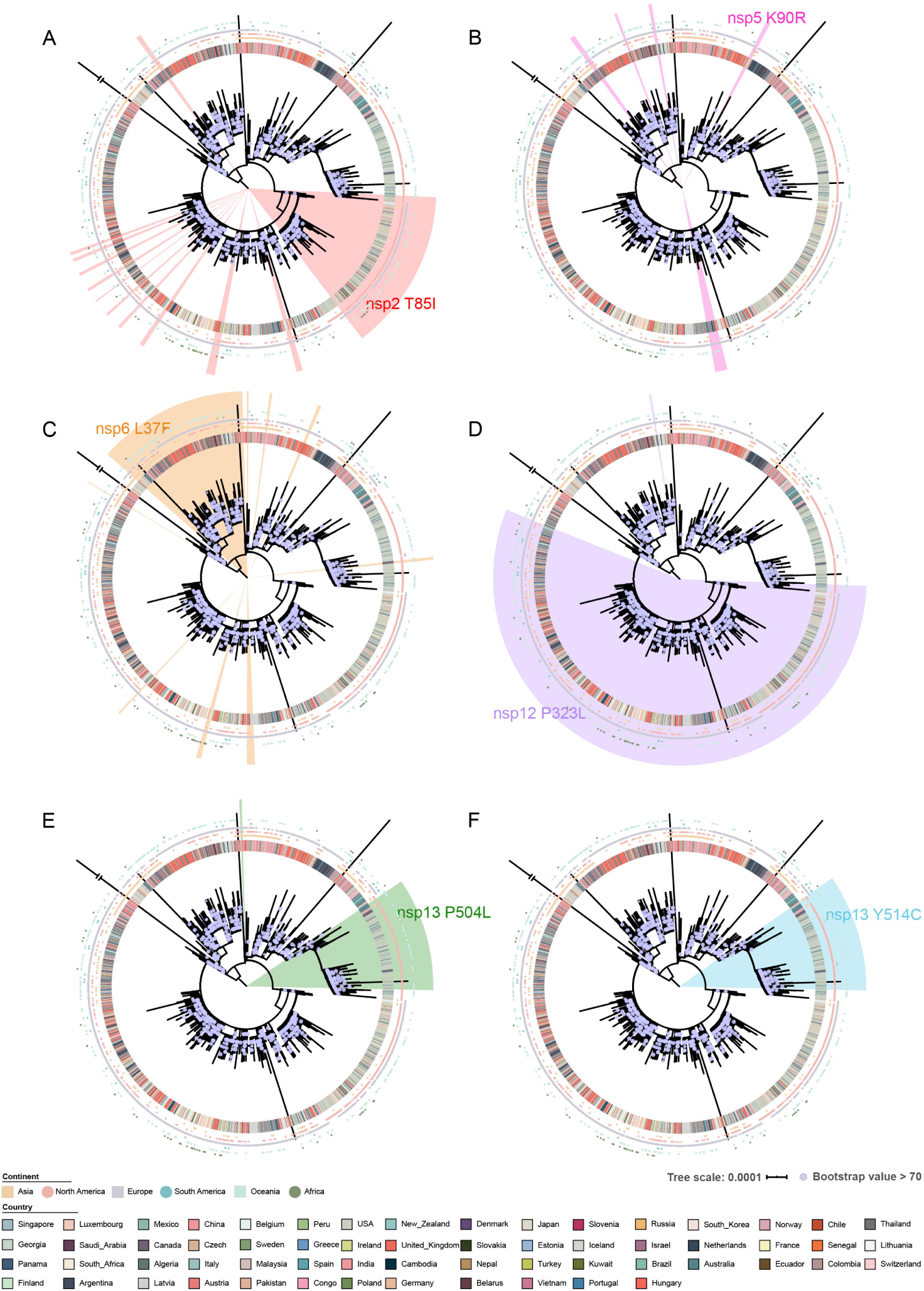
Phylogenetic tree of ORF1ab including the location of putatively positively selected sites and sites exchanged in more than 10% of SARS-CoV-2 sequences. The outer circle represents the origin of a sequence from a certain continent, and the inner circle from a certain country, as specified at the bottom. The triangle-shaped shadow represents the position of the mutant strains in the phylogenetic tree. Different colors represent different mutation sites. **A**: T85I in nsp2, a site, with a mutation frequency of 15.47%. **B**: K90R in nsp5 are sites dN/dS > 1, but occur only in more than 1% of sequences. **C**: L37F in nsp6, a site with a mutation frequency of 14.02%. **D**: P323L in RdRp in nsp12, a site with a mutation frequency of 56.10%. **E**: P504L in nsp13 a site, with a mutation frequency of 10.08%. **F**: Y541C in nsp13 a site, with a mutation frequency of 10.38%.

Since nonsynonymous changes are more likely to associate with functional adaptations of viruses, we searched for amino acids substitutions in our collection of 4,894 whole genome sequences. We identified 11 amino acids which are mutated with high frequency (more than 10%) in SARS-CoV-2 sequences. Two of the sites are located on ORF1a (nsp2, nsp6), three on ORF1ab (nsp12, nsp13), one on S, two on ORF3a, one on ORF8 and two on N (Table 1). On the ORF1ab phylogenetic tree, five high-frequency mutant strains show branch aggregation and one site (nsp5 Lyr90Arg, mutation frequency more than 1%) shows a scattered distribution (Fig. 2). Whereas four of the five high-frequency mutant strains can almost cover one cluster and the other (nsp12 Pro323Leu) is very widely distributed, covering almost three clusters. Note that the two mutations in nsp13 are clustered together and occurred mainly in North America, the other mutations are present in different clusters of various geographic origin.

**Table 1.** Frequently exchanged amino acids in SARS-COV-2 genomes.

| Gene | Protein | Position | Nucleotide mutation | Amino acid mutation | Mutation frequency | Geographic Sample and Major Country |
| --- | --- | --- | --- | --- | --- | --- |
| ORF1a | nsp2 | 85<sup>#</sup> | ACC→ATC | Thr→Ile | 15.47% | 23 Country, USA, IS, FR, AU, NL |
|  | nsp6 | 37<sup>#</sup> | TTG→TTT | Leu→Phe | 14.02% | 36 Country, UK, AU, JP, NL, USA |
| ORF1ab | nsp12, RdRp | 323 | CCT→CTT | Pro→Leu | 56.10% | 49 Country, USA, UK, NL, IS, BE |
|  | nsp13 | 504 | CCT→CTT | Pro→Leu | 10.08% | 5 Country, USA, AU, CA, IS, UK |
|  |  | 541 | TAT/TAC→TGT | Tyr→Cys | 10.38% | 5 Country, USA, AU, CA, IS, UK |
| S | S1 | 614<sup>#</sup> | GAT→GGT | Asp→Gly | 56.49% | 49 Country, USA, UK, NL, IS, BE |
| ORF3a |  | 57 | CAG→CAT | Gln→His | 19.06% | 28 Country, USA, FR, IS, AU, UK |
|  | ORF3a | 251 | GGT→GTT | Gly→Val | 10.25% | 27 Country, UK, AU, NL, CH, IS |
| ORF8 | ORF8 | 84 | TTA→TCA | Leu→Ser | 16.13% | 27 Country, USA, CH, AU, CA, ES |
| N | N | 203 | AGG→AAA | Arg→Lys | 16.30% | 27 Country, UK, NL, BE, IS, AU |
|  |  | 204 | GGA→CGA | Gly→Arg | 16.27% | 40 Country, UK, NL, BE, IS, AU |

Amino acids with a mutation frequency greater than 10%.
“#” indicates that the site might be a positively selected site (see table S3)

Figure 3 shows a more detailed temporal and spatial analysis: the percentage of a certain mutation in all sampled sequences and in sequences from Asia, Europe and North America is plotted against the time period when the sequences were deposited.

**Figure 3:**
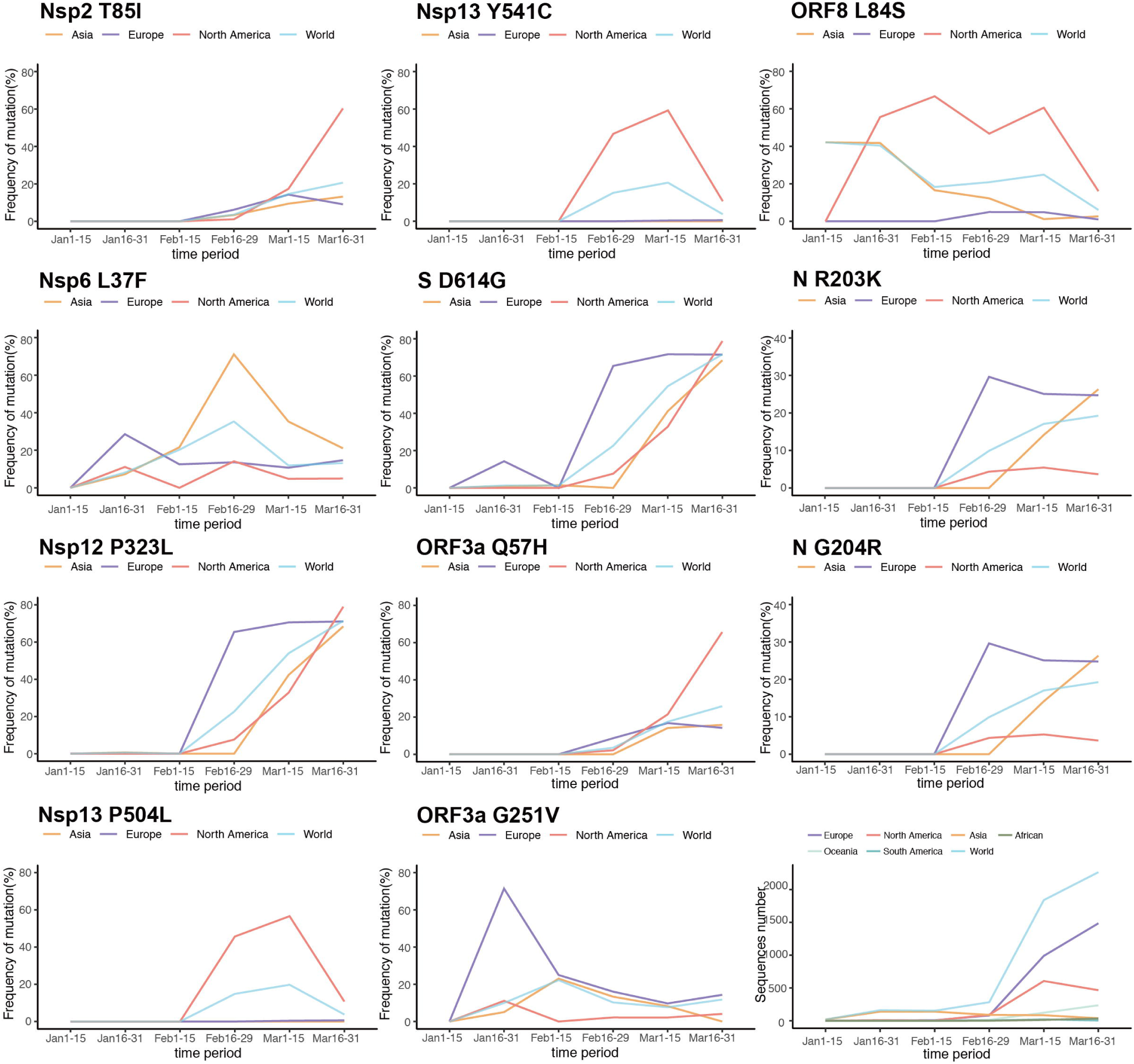
Temporal and spatial analysis of high-frequency mutation sites. The frequency of a certain mutation in three continents (Asia, Europe, North America) and in all sequences (world) is plotted against the time period these sequences were submitted to the database. The graph at the bottom right shows the number of submitted nucleotide sequences for each time period.

Some mutant viruses appeared with a high but declining frequency in January in certain regions that presented pronounced local heterogeneity. For example, ORF3a Gly251Val was highly abundant in Europe in the second half of January (71% versus 5-11% in Asia and North America), whereas the mutation ORF8 Leu84Ser was very frequent in North America and in Asia in the second half of January (40-60%), but absent in Europe (0%). The mutation in nsp6 revealed a more fluctuating pattern with local variation (71% in Asia, but only 14% in Europe and North America), but this isolate is now of low abundance throughout the world.

Other mutations occurred first in February with declining frequencies and notable local heterogeneity. Both mutations in nsp13 appeared in North America in the second half of February at high frequency (∼46%), which increased to >55% and then dropped sharply to 11%. These mutations in nsp13 were observed only at very low frequency in Europe (<1%) and not at all in Asia.

Interestingly, some mutations presented an increasing abundance over time. For example, the mutations in S and nsp12 are highly abundant in all regions (>68%) in the second half of March as also observed in another study published very recently^32^. An increase in the frequency of the mutation in NSP12 and S first occurred in China in the second half of January. The mutations in ORF3a Gln57His and nsp2 Thr85Ile reached a high frequency in North America (>60%) in March but remained lower in Europe and Asia (9-16%). Both mutations in the N-protein are also increasing in frequency, first in Europe (up to 30% already in February) and more recently in Asia (26% in March).

Note that the graphs for S and nsp12 are similar to each other in all regions, and to the graphs of both mutations in the N-protein. When analyzing individual sequences, we found that the two mutations in the N-protein as well as the mutations in S and nsp12 very often appeared together in one genome. The strong association of the mutation sites S D614G and nsp12 P323L (together with a C-to-T mutation at position 241 of the 5’ UTR and a silent C-to-T nucleotide exchange at position 3,037) has been described also in another study published very recently^32^. We also observed quadruple mutations: viruses with the mutation in S and nsp12 acquired the two mutations in the N gene, but the additional mutations were not always retained. The same pattern is consistent for the mutations T85I in nsp2 and Q57H in ORF3a that often occurred together in one genome and also as a quadruple mutation together with the mutations in S and nsp12. Thus, some of the mutations are inherited together explaining their similar local and spatial occurrence.

There are 17 additional amino acid exchanges if we lower the threshold value (>1%), present in all genes, except for the genes encoding E, ORF7a, ORF7b, and ORF10. The additional mutation sites are mostly located in ORF1ab with one in ORF2, one in ORF8 and two in the M and N genes (Table S2). Of note, except for the highly abundant Asp614Gly exchange, there were no further changes identified in the spike protein. Further tracking the occurrence of these mutations in the ongoing pandemic might provide additional insights into their roles.

In addition, we found that ten of the eleven high-frequency amino acid substitutions were caused by a one nucleotide change, 7x at the second position, 2x at the third position and 1x at the first position of a codon. Only one mutant exhibits an exchange of two nucleotides: Arg encoded by AGG changed to AAA specifying Lys at amino acid position 203 of the N gene. Since the viral polymerase makes at most one nucleotide substitution per replication cycle, a double nucleotide exchange is unusual.

In all 12 mutation sites the variable residue was always exchanged to the same amino acid, although more replacements would be possible by a one nucleotide exchange. Only four of the amino acid mutations are conservative replacements, such as the exchange of the basic residues Arg and Lys and of the hydrophobic residues Leu to Phe. Five exchanges involve the unusual amino acids Gly and Pro, which destabilize secondary structures, and are not able to interact with other amino acids.

In order to explore a potential adaptation of SARS-CoV-2 to humans, we estimated dN/dS ratios for the ORF1ab and S protein phylogenetic trees using four algorithms (Fig. 2 and Fig. S2). Interestingly, some high-frequency mutations exhibit dN/dS >1: Thr85Ile in nsp2, Leu37Phe in nsp6, Asp614Gly in S and Pro323Leu in nsp12. Note that the frequency of three of these sites is increasing in almost all continents since their first appearance, and that only Leu37Phe in nsp6 is decreasing in frequency (Fig. 3). Another site, Lys90Arg in nsp5 was also identified by the dN/dS analysis, which has a mutation frequency higher than 1% (Fig. 2b). Thus, this suggests that a certain number of sites within SARS-CoV-2 have potentially undergone positive selection. However, the increase in the frequency may coincide with the transmission pattern of the virus in different regions and further analysis is needed to substantiate this preliminary conclusion.

### 2. Structural analysis of frequently exchanged sites in non-structural proteins

**Nsp2**, a protein that modulates host responses, contains one site with exchanges greater than 15%, the exchange Thr85Leu, the site might be positively selected, but no structural information is available for nsp2 of any β-coronavirus to specify any function for this site.

One high-frequency site for which some evidence of selection is available can be found in **nsp6**, a membrane protein with six transmembrane spans which induces the formation of autophagosomes from the ER membrane^33^. The mutation we found (Leu37Phe) is located at the end of the first transmembrane region directed to the lumen of the ER. The physiological role of this exchange is obscure since residue 37 is already preceded by three Phe, i.e. the amino acid sequence changes from LFFF<u>L</u>Y to LFFF<u>F</u>Y.

One site with mutation frequency greater than 50% was identified in **nsp12**, the RNA-dependent RNA polymerase (RdRp)^7^. Nsp12 forms a complex with one copy of nsp7 and two copies of nsp8 (Fig. 4a). It is divided into the catalytic part common to all RNA-viruses (green colours) containing the canonical “palm”, “finger” and “thumb” domains and N-terminal extension unique to Nidoviruses, which is composed of a domain with nucleotide transferase activity (NiRan, magenta) and an interface domain (blue)^34^. The selected site, residue 323, is located in the interface domain at the surface of a cavity at the backside of the polymerase complex. The back wall of the cavity is built by the “finger” domain, its left and right part are formed by nsp8 and the interface region, respectively (Fig. 4b). The opening of the cavity is flanked by two basic amino acids and its back contains several hydrophobic residues (Fig. 4c). Proline 323 is located in a short, twisted α-helix; its side chain is exposed at the surface of the cavity (Fig. 4d). Exchanging proline with the hydrophobic leucine is increasing the hydrophobic character of the cavity.

**Figure 4:**
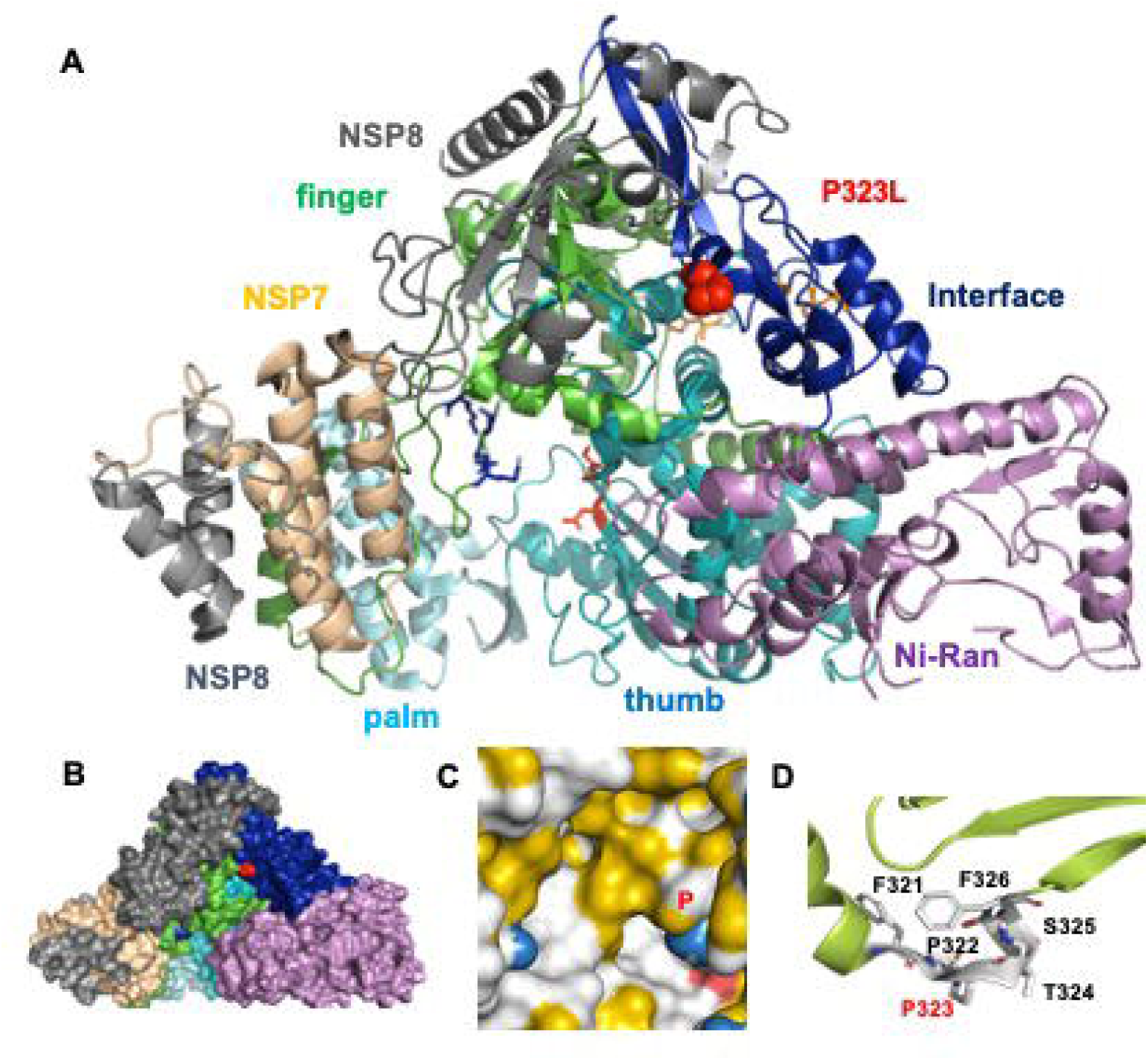
Amino acid frequently exchanged in the polymerase complex. **A:** Crystal structure of the nsp12 polymerase in complex with one copy of nsp7 (coloured wheat) and two copies of nsp8 (grey). The NiRan domain of nsp12 is coloured in magenta and the interface region in blue. The location of residue 322 is shown as red sphere. The catalytic domain is coloured from green to light blue indicating the palm, finger and thumb domains. The catalytic amino acids are shown as red sticks. **B:** Surface representation of the same complex. Pro 322 is coloured red. Colouring of the domains is as in A. **C:** Hydrophobic cavity with the location of Pro 322 (labelled P). Three hydrophobic surface patches in the back are in yellow (from left to right: V398, A399; V535, I536; V320, F321) and two basic amino acids (K391, R654) at the entrance are in blue. **D:** Detail of the structure showing the location of Pro 323 and adjacent amino acids as sticks. Figures were created with PyMol from pdb file 6M71^34^. Inspection of the other available nsp12 structures either bound to remdesivir (pdb 7BV2, ^54^) or of a replicating SARS-CoV-2 polymerase (pdb 6YYT, unpublished) revealed the same results.

Two identified high-frequency mutation sites were located on **nsp13**, a helicase that unwinds DNA or RNA duplexes^8^. We used the structure of the highly similar nsp13 of MERS to highlight the location of the residues^35^. Nsp13 is composed of a N-terminal CH domain, which is connected by a stalk domain to two RecA domains. The two exchanged residues, proline 504 and tyrosine 541, are located at the surface of the RecA2 domain (Fig. 5a). The side chain of Tyr 541 runs parallel to the surface and forms a hydrogen bond with Gln531, which is lost if substituted by a cysteine. The main chain of Pro 504 interacts with Ser507. This interaction is preserved when Pro504 is exchanged by a Leu, but the substitution exposes a hydrophobic moiety at the surface of NSP13. Thus, both exchanges might alter the surface properties of nsp13, perhaps to modify binding to other viral or cellular factors.

**Figure 5:**
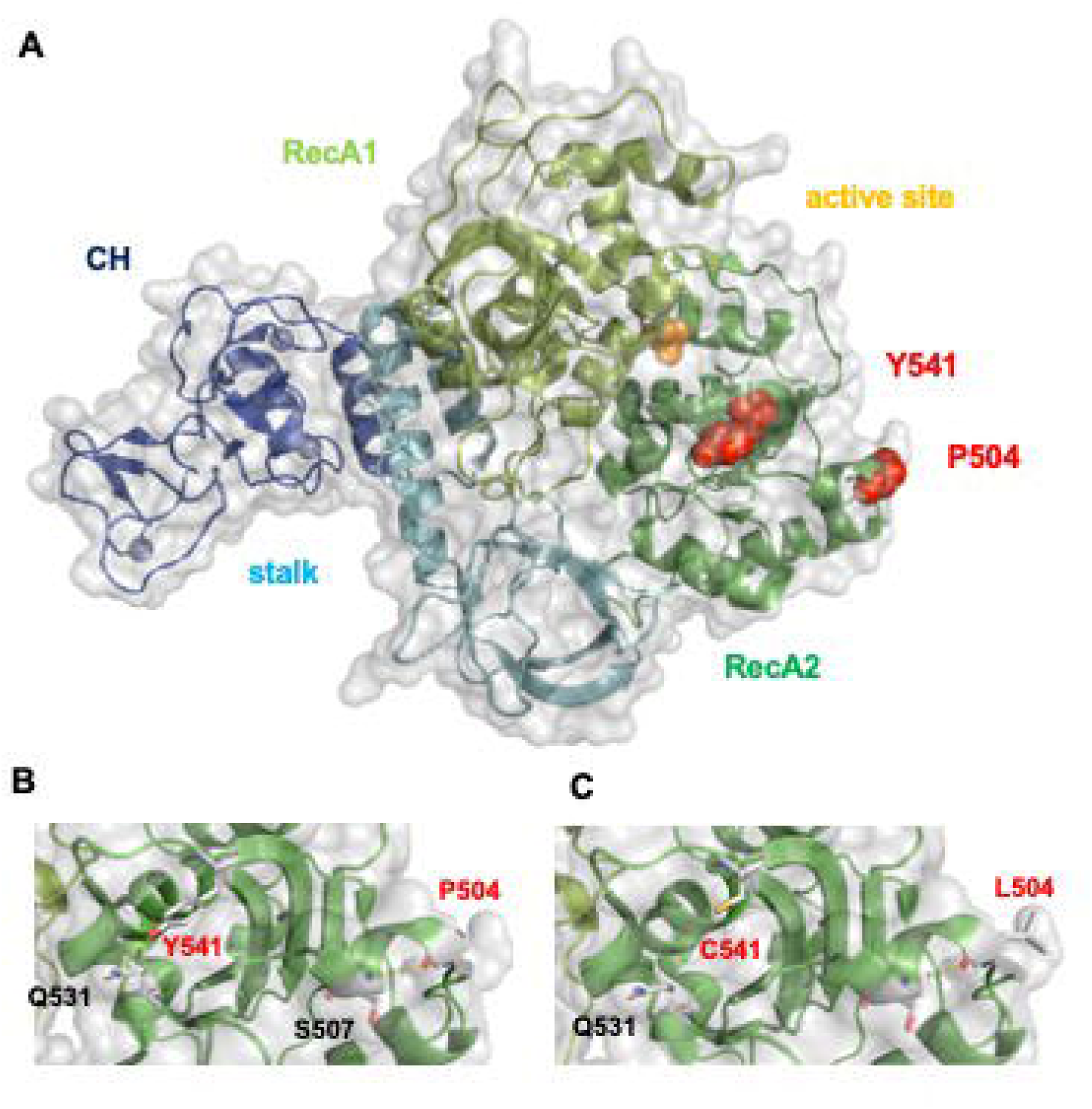
Amino acids frequently exchanged in nsp13. **A**: Structure of nsp13 from MERS has 72% identical and 85% similar amino acids compared to nsp13 of SARS-CoV-2. The individual domains are coloured and labelled, CH domain in blue, the stalk in light blue, the two RecA domains in light and dark green. The active site contains a sulfate (yellow sphere). Pro504 and Tyr541 are shown as red spheres. **B**: Surface representation of nsp13 with Pro504 and Tyr541 and their interaction partners indicated as white sticks. The main chain of Pro504 interacts with Ser507, whereas the side chain of Tyr541 forms a hydrogen bond with Gln 531. Note that position 507 is an Arg in nsp13 of SARS-CoV-2, but this exchange is unlikely to disturb the interactions with the main chain of Pro504. **C**: Exchange of Tyr541 with a cysteine destroys the hydrogen bond whereas substitution of Pro504 by Leu exposes a hydrophobic side chain at the molecule’s surface. All figures were created from pdb file 5WWP^35^. Homology modelling of nsp13 from SARS-CoV-2 using swiss model produced the same results.

One site which is possibly positively selected, but exhibits a low mutation frequency, is located in **nsp5**, the main protease of coronaviruses. Nsp5 exists as a dimer and dimer formation is essential for catalytic activity^36^. Fig. 6c shows the structure of an nsp5 monomer from SARS-CoV-2 with the active center (His41, Cys145) shown in red and other residues involved in substrate binding as orange sticks^34^. They are located in domain II which is connected by a long loop to domain III which contains amino acids important for dimerization (shown as blue sticks). The selected site, lysine 90, is located in domain I at the other side of the molecule. The arginine is part of a ß-sheet, its side chain is exposed at the surface of the molecule where it forms a hydrogen bond with Asp34 (Fig. 6d). The exchange of Lys90 by the other basic amino acid Arg does not change the electrostatic properties of the molecule’s surface, but the hydrogen bond to Asp34 might be lost, since the side chain of Arg is longer (Fig. 6e).

**Figure 6:**
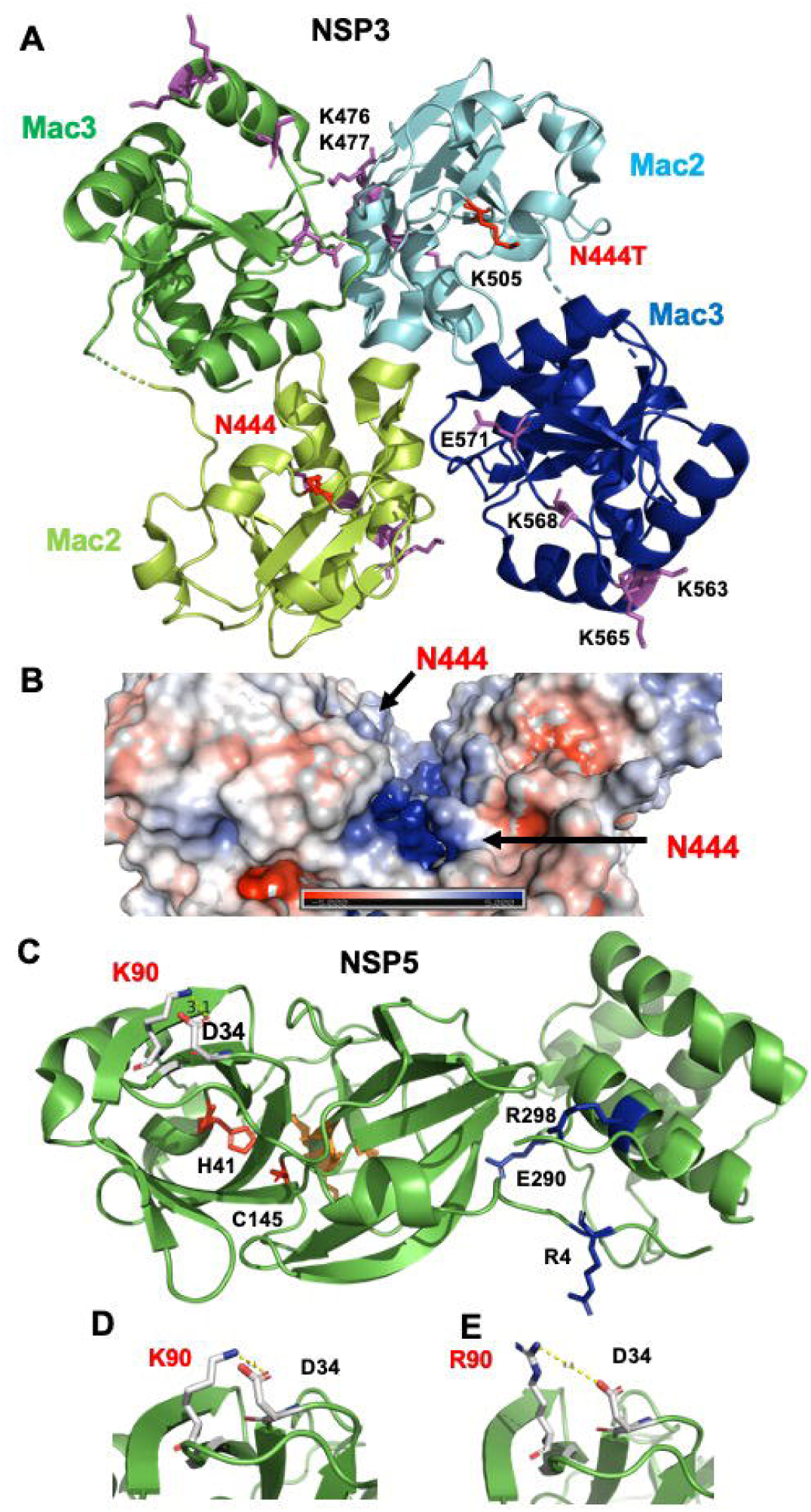
Amino acid under positive selection in nsp3 and nsp5. **A:** Dimer of mac 2 and mac 3 domains of nsp3 of SARS-CoV-1, which exhibits 75% identical and 86% similar amino acids compared to the corresponding polypeptides of SARS-CoV-2. Domains which belong to one monomer are labelled blue or green. The location of the selected amino acid Asn 444 (which is a Lys at position 420 in SARS-CoV-1) is shown as red stick. Basic amino acids which are involved in poly(G) binding are shown as magenta sticks and their number is indicated in one monomer. **B:** Electrostatic surface potential of a groove between monomers that runs perpendicular to the poly(G) binding cleft. Asn 444 is located at the entrance of the cleft. The bottom of the cleft contains the basic amino acids Lys462, Lys463 and Arg473. Nsp3 from SARS-CoV-2 also has basic amino acids at the positions mentioned in A and B. Figures were created with Pymol from pdb file 2W2G^60^. Homology modelling of nsp3 from SARS-CoV-2 using swiss model (https://swissmodel.expasy.org/) produced the same results. **C:** Crystal structure of a nsp5 monomer. The catalytic residues (His41, Cys145) are shown as red sticks and residues involved in substrate binding (His163, Met165, Glu166, His172) as orange sticks. Amino acids important for dimerization (Arg4, Glu 290, and Arg298) are shown as blue sticks. The selected site, Lys 90, is shown as white stick. **D:** Detail of the structure showing the location of Lys90. Its side chain is exposed at the surface of the molecule where it forms salt bridge (distance 3.1Å) with Glu34. Created with Pymol from pdb file 6Y2E^34^. Another available structure of nsp5 from SARS-CoV-2 (pdb file 7BQY ^34^) also revealed a hydrogen bond between Lys90 with Glu 34. **E:** Exchange of Lys90 by Arg90 increases the distance to Glu34 to 4.8Å.

### 3. Structural analysis of frequently exchanged sites in accessory proteins and virus components

Two frequent exchanges were identified in **ORF3a**, a homotetrameric ion channel of unknown structure that also activates inflammatory responses of the host^37,38^. The first exchange Gln57His is located at the cytosolic end of the first transmembrane region and the second exchange Gly251Val in the cytoplasmic region.

**ORF8**, a small glycoprotein of unknown function, contains one frequently exchanged site at position 84, a hydrophilic serine is exchanged by hydrophobic leucine. Note that ORF8 often acquires mutations after the host jump of a SARS virus. A 382-nucleotide deletion covering almost the entire ORF8 was reported for SARS-CoV-2 isolates from eight hospitalized patients^39^. Likewise, for ORF8 of SARS-CoV-1 it has been reported that a 23-nucleotide deletion acquired early during human-to-human transmission attenuates virus replication in cell culture^40^.

Two sites were identified in the **N protein**, which packages the viral RNA into ribonuleoparticles. N consists of a N-terminal RNA-binding domain and a C-terminal oligomerization domain, which are connected by an unstructured linker region that harbours the two penultimate exchanges^41^. It contains an Arg/Ser rich region with various phosphorylation sites, but the exchanges do not change the probability for phosphorylation of any of the sites.

The S protein contains one selected exchange (Asp614Gly), which is located in the S1 subunit between the receptor-binding domain (RBD) and the furin cleavage site (Fig 6a). S2 harbours the fusion machinery of the viral spike: a hydrophobic fusion peptide adjacent to the second proteolytic cleavage site (S2) and a heptad repeat regions (HR) (Fig. 7b). Asp614 forms the apex of a loop that links a short β-strand to an unresolved, hence probably flexible part of S1. The loop contains a used N-glycosylation site, Asn 616. It is covalently connected to another loop by a disulphide-linkage between Cys 617 and Cys 649 and via interaction of the main chain atoms of Asp614 with Ala647. Both loops are at the interface between two monomers of the trimeric spike protein. Interestingly, the carboxyl group of Asp 614 forms a hydrogen bond with the -OH group of Thr859, which is part of a short β-strand in the S2 subunit between the fusion peptide and the heptad region 1 and is (Fig. 7b). S of SARS-CoV-1 also exhibits a similar interaction between monomers, Asp 600 forms a hydrogen bond with Thr841 and the other elements characteristic for this region, are also present (Fig. 7c).

**Figure 7:**
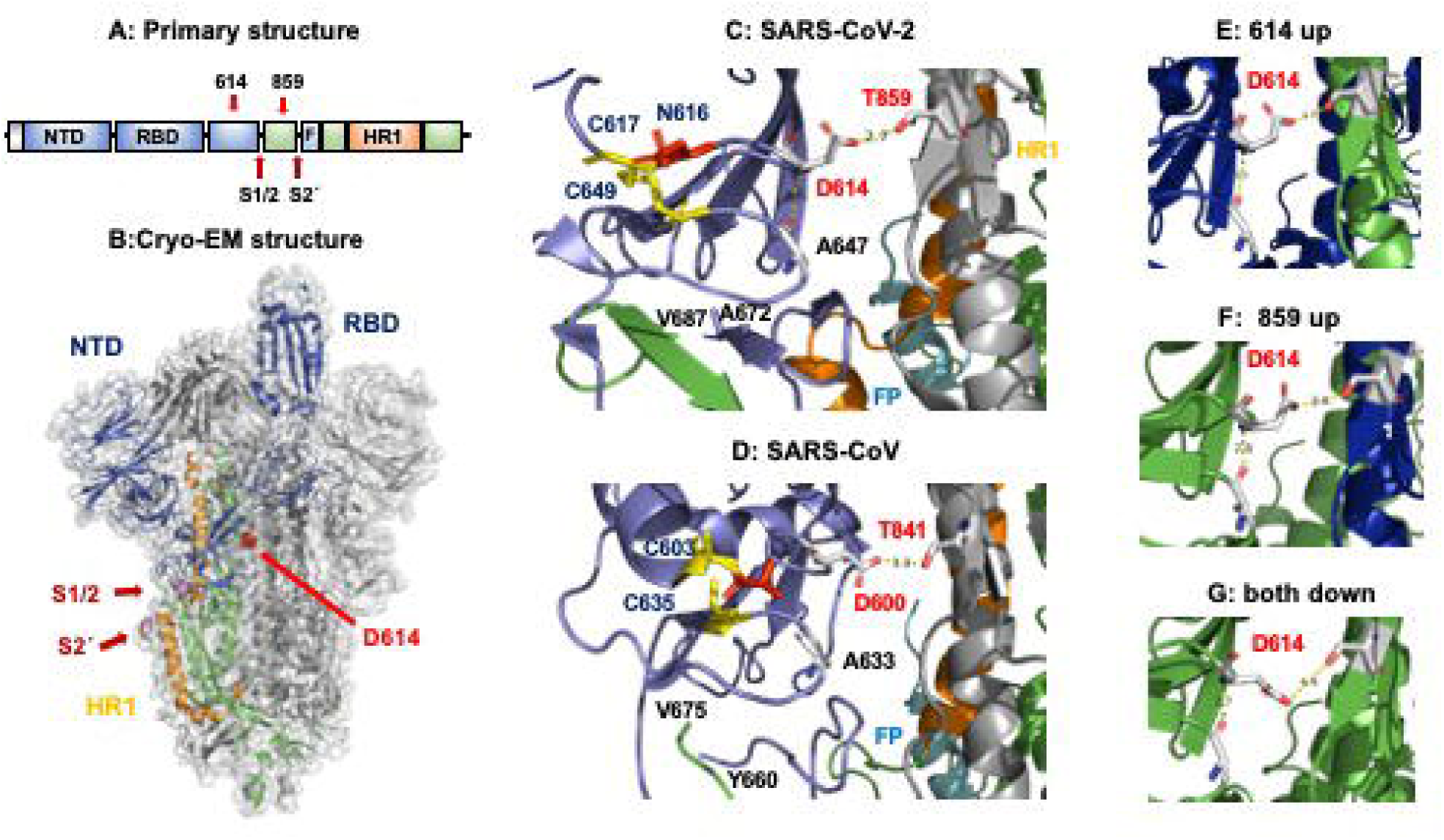
Amino acid frequently exchanged in the spike protein. **A:** Primary structure of S showing the location of positions 614 and 859. The S1 subunit is in blue and contains the N-terminal domain (NTD) and the receptor binding domain (RBD). The S2 subunit is in green except for the heptad repeat region 1 (HR1), which is in orange and the fusion peptide F, which is in blue. The proteolytic cleavage sites S1/2 and S2’ are indicated by red arrows. **B:** Location of D614 in the trimeric Cryo-EM structure of S. Protein domains in one monomer are coloured as in A. The RBD of the coloured monomer is in the up conformation, the other two in the down conformation. Pdb file: 6VSB^61^) **C:** D614 in S1 forms a hydrogen bond with Thr859 in S2 of another monomer. Protein domains are coloured as in A. Pdb file: 6VSB. **D:** Uncleaved S of SARS-CoV-1 with all RBDs in the down conformation: D614 in S1 forms a hydrogen bond with Thr841 in S2 of another monomer (pdb file 5WRG^4^). Protein domains are coloured as in A. **E**: Distance between D614 in the up monomer and Thr 859 in the down monomer. **F**: Distance between D614 in the down monomer and Thr 859 in the up monomer. **G**: Distance between D614 and Thr 859 both in the down monomer. E to G were created with PyMol from pdb file 6VSB. Note that in the other available Cryo-EM structures of S (pdb files 6VXX and 6VYB ^62^) the distance between Glu614 and Thr 859 is larger. However, since not all side chains are properly displayed in PyMol, this might be due to a lower resolution in this region.

Individual monomers of the prefusion structure of S exists in several conformations: The RBD could be either hidden from receptor engagement (“down” conformation) or exposed by a hinge-like movement at the top of the molecule (“up”) that would allow receptor-binding. Interestingly, the distance between the carboxyl group of Asp and the -OH group of threonine is the shortest (2.7Å) if Asp is a component of an up-monomer (Fig. 7d). The distance becomes larger (3.4Å) if Thr is present in an up monomer (Fig. 7e) or if both monomers are in the down conformation (4.4Å) (Fig. 7f). Analysis of resolved structures of S of SARS-CoV-1 showed that binding to the ACE2 receptor or to antibodies, which both move the RBD into the up conformation, increased the distance between Asp and Thr to more than 5 Å^42,43^, which is too large to maintain the hydrogen bond between Asp and Thr^44^. This suggests that exposing the RBD at the top of the molecule is accompanied by small movements of the S1 and S2 subunits against each other. The exchange of Asp614 by a glycine would prevent hydrogen bond formation and thus might facilitate exposure of the RBD.

In the S-protein, we also searched for sites which are exchanged in only ten or more sequences. However, none of the amino acids making up the epitopes of neutralizing monoclonal antibodies generated from memory B-cells of human SARS-CoV-1 or SARS-CoV-2 patients were detected^45,46^. Likewise, there was just one mutation in the whole receptor binding domain, residue 483 changed from Val to Ala in 16 sequences. However, this residue does not directly contact the ACE receptor^47^. Thus, there is no evidence that SARS-CoV-2 can easily generate antibody escape mutants or is adapting to recognize the human receptor more efficiently.

## Discussion

The evolution and epidemiology of a new virus at different stages after its spread into a new host are often considered to be critical for its success in creating and sustaining an epidemic. As of 20th June 2020 more than 49,000 complete or near-complete genomic sequences of SARS-CoV-2 are publicly available and the number rises daily. These genomes provide invaluable insights into the ongoing evolution of the virus during the pandemic, which might be helpful to eventually mitigate and control the virus spread. Until now, there is no official definition of the SARS-CoV-2 phylogenetic lineages, but a few articles have proposed methods of classification. Although the classification methods are diverse, the method of Rambaut et al [52] is the current mainstream which clearly defines the principles of lineage classification. We reconstructed phylogenetic trees based on early 4,894 full genomes, ORF1ab and S gene sequences, which is by now the one of the most comprehensive analysis of the early epidemic/mutations characteristic of SARS-COV-2. Using full genomes we identified five major clusters in the phylogenetic tree which is similar to other observations (https://virological.org/t/year-letter-genetic-clade-naming-for-sars-cov-2-on-nextstain-org/498), but it is too early to denote new genotypes of SARS-COV-2, because sequences are highly homogeneous and the phylogenetic trees exhibit low bootstraps values and no significant changes in clinical symptoms or transmissibility have been reported so far. Although some manuscripts report that SARS-COV-2 has produced new genotypes^18-20^, our phylogenetic analysis shows that the genetic diversity of SARS-COV-2 is relatively low during this early stage of the epidemic, the viral genome is largely stable and the virus did not evolve rapidly after its emergence in humans. However, each of the five clusters has its own mutations that may serve as targets for fast genotyping of samples from patients. Our nomenclature may complement the system suggested by Rambaut et al^48^ which proposed a dynamic system for labeling transient lineages that have local epidemiological significance.

In addition, on the amino acid level we identified eleven mutations that occur in more than 10%, two of them even in more than 50% of the analyzed SARS-CoV-2 sequences (Table 1). Although some mutations have been reported before^49-52^, most of the high frequency mutations in ORF1ab are new findings.

We analyzed these mutations further by using four algorithms to calculate dN/dS for each site. We found that four of them exhibit a dN/dS larger than 1, three are in ORF1ab (Thr85Ile in nsp2, Leu37Phe in nsp6, Pro323Leu in nsp12) and one in the S protein (Asp641Gly) (Table S3). Other sites also exhibit a dN/dS value greater than 1, but the frequency of these exchanges is currently very low. Although these sites have selection signals, the selection analysis has some limitations and hence we consider the results as preliminary. It has been reported that some mutations that seemingly arise multiple times along the phylogenetic tree may be caused by sequencing error and/or are the result of either artefactual lab recombination, or potential hypermutation^53^ (https://virological.org/t/issues-with-sars-cov-2-sequencing-data/473). In addition, during an ongoing pandemic, purifying selection signals occur frequently and recombination events between viruses might obscure the signal^53^. Furthermore, we cannot exclude that some of the exchanges are the result of non-selective conditions, i.e. due to a “genetic bottleneck”. A patient releases droplet that contain only a limited virus population which does not representative of the whole virus “swarm” replicating in his body. One droplet that by chance does not contain a single particle representing the original strain might then infect another person where it creates a new and different virus population by a ‘founder’ effect. However, since the pandemic continues and new viruses will be sequenced, it is worthwhile to analyze whether one of the sites become positively selected.

In summary, our study revealed that in the early large genome of SARS-CoV-2 only a few amino acids are exchanged and hence the selection pressure is low, which is consistent with the conclusion in the good sequence analysis reported by MacLean et al ^53^.

We also performed a spatial and temporal analysis that revealed the time and region at which a certain mutation appeared first and how it spread to other continents together with its frequency change in the sampled population. The mutations in nsp6, nsp13, ORF3a 251 and ORF8 exhibit high-frequency peaks early in the epidemic and only in certain regions, but then showed a sharp drop and are of low abundance now. In contrast, the frequency of the mutations in nsp2, nsp12, ORF3a 57, N and S increased since their introduction into the viral genome. They are present at high abundance in the second half of March, either worldwide (S, nsp12, >68%) or in certain continents (N in Europe and Asia, ∼25%, nsp2 and ORF3a in North America, >60%).

For the viral proteins where 3D structural information is available, we performed an in-depth analysis of possible consequences of these amino acid exchanges. One conclusion is that none of the exchanges is presumed to disturb the folding of the protein. The selected amino acids are located either in loops (Asp614 in S) that are flexible and could probably accommodate many amino acids or at the edge of β-sheets (Lys90 in nsp5, Tyr451 in nsp13).

Another similarity between the identified sites is that they are located at or very close to the surface of the molecule. The crystal structures revealed that the side chains of three of the sites interact with other amino acid in close proximity. Lys90 in nsp5 forms a salt bridge with Asp34, whereas Tyr 541 in nsp13 and Asp614 in S form hydrogen bonds with Gly531 and Thr859, respectively. In all three cases the interaction is destroyed by the substituted amino acid.

Two other frequently exchanged residues are prolines, which do not have a side chain that can interact with other amino acids. They are substituted in nsp12 and nsp13 by a leucine residue, which then exposes a new hydrophobic moiety at the molecule’s surface. Especially interesting is the exchange in nsp12, the RNA-dependent RNA polymerase, since it is potentially positively selected and occurs in more than 50% of SARS-CoV-2 sequences. Pro 323 is located on the back side of a hydrophobic cavity, its exchange by a leucine increases its hydrophobic character.

In conclusion, the exchanges in nsp2, nsp5, nsp13 and S alter the local biophysical properties at the molecule’s surface and expose new amino acids, which could potentially interact with other molecules. One might thus speculate that the identified residues are elements of an interaction surface with cellular polypeptides, e. g. one of the 323 human proteins that have been identified to interact with SARS-CoV-2 proteins^54^. Since the virus has been only recently introduced into the human population, likely from bats via an intermediate host ^2^, it is reasonable to assume that they are not perfectly adapted to human cells. The mutations thus might increase the affinity for a human polypeptide that differs in the interaction surface from the ortholog protein from the previous hosts of SARS-CoV-2. The physiological role might be to better antagonize the innate immune response of the host, but the Coronavirus proteins that fulfil this tasks have not been carefully explored ^55^. In addition, improved interaction with cellular proteins might facilitate gene expression (nsp5, nsp13), RNA processing (N), signaling (nsp13, N) and especially vesicle trafficking (nsp2, nsp6, nsp13, ORF3a, ORF8), the latter perhaps to facilitate the reconfiguration of ER/Golgi membranes to create the replication and transcription complex^54^.

Bat viruses are also adapted to grow at a wide range of temperatures, ambient temperature during sleeping and up to 41°C during flight ^56^. Thus, proteins from bat viruses must withstand higher temperatures, which occur in humans only as part of the antiviral defense mechanism. Thermostability of proteins arises from the simultaneous effect of the forces between its amino acids^57^. Interestingly, two of the exchanges we identified (Tyr541Cys in nsp13 and Asp614Gly in S) likely cause a loss of a salt bridge or hydrogen bond, whereas the opposite effect, a gain of a new interaction by the substituted amino acid could not be predicted. The loss of bonds increases the local flexibility of a polypeptide chain, which might enhance protein-protein interactions ^58^.

A more specific speculation can be offered for the potentially positively selected and high frequency site in the spike protein S. Asp614 in the S1 subunit forms a hydrogen bond with Thr859 in the S2 subunit of an adjacent monomer. The substitution to a glycine abolishes the hydrogen bond and thus might destabilize the trimeric spike protein. This in turn might affect exposure of the receptor-binding domain at the top of the molecule, which is a prerequisite for binding to ACE2. The mutation might also affect the fusion activity of S, which requires dissociation of the S1 subunit from the spike before the canonical conformational change of S2 can occur that executes membrane fusion. The trigger for the conformational change is unknown, but it seems possible that a slightly destabilized trimer might be more prone to refolding compared to S having the Asp residue at position 614.

Functional consequences of the Asp614Gly exchange have recently been published. Two pseudoviruses having a Gly at position 614 grow to 2-9-fold higher titers in various cell types compared to pseudoviruses having the Asp. In addition, COVID-19 patients infected with a virus with the G614 mutation tend to exhibit higher virus load, but no association with disease severity was found. Furthermore, both pseudoviruses are neutralized with equal efficiency with convalescent sera suggesting that this locus is not critical for antibody-mediated immunity^32^. Therefore, although S Gly614 has a unique phenotype, more research is needed to decide whether it increases virus entry and/or transmissibility between humans.

Here, by studying 4,894 sequences from an early stage of the epidemic, we have created an analysis pipeline to track potential adaptive mutations in SARS-CoV-2. Whether any of the identified exchanges alter clinical properties of SARS-CoV-2, such as transmissibility, cell tropism, replication rate and pathogenicity needs further explorations. Usually, such important traits are controlled by multiple genes ^59^. Since essentially no exchanges occurred in the receptor binding domain of the spike protein and in the epitopes recognized by neutralizing antibodies, there is no evidence that SARS-CoV-2 adapts to improve receptor binding and/or to escape from the adaptive immune response. Nevertheless, it seems appropriate to follow further the evolution of SARS-CoV-2 during the ongoing pandemic.

## Acknowledgements

SS, WTH, JZ, JMS, XFZ are financially supported by the National Key Research and Development Program of China (Grant no. 2016YFD0500402), National Ten Thousand Talent Program, the Fundamental Research Funds for the Central Universities (Grant no. Y0201900459), Six Talent Peaks Project of Jiangsu Province of China (Grant no. NY-045) and the Bioinformatics Center of Nanjing Agricultural University. XJ and MAS are partially supported through National Institutes of Health grant U19 AI135995. GB acknowledges support from the Interne Fondsen KU Leuven / Internal Funds KU Leuven under grant agreement C14/18/094, and the Research Foundation – Flanders (‘Fonds voor Wetenschappelijk Onderzoek – Vlaanderen’, G0E1420N).

## Tables

**Table S1.** Between group mean p-distance of five clusters in whole genome phylogenetic tree.

|  | Cluster I | Cluster II | Cluster III | Cluster IV | Cluster V |
| --- | --- | --- | --- | --- | --- |
| Cluster I |  |  |  |  |  |
| Cluster II | 1.61E-04 |  |  |  |  |
| Cluster III | 1.04E-04 | 1.53E-04 |  |  |  |
| Cluster IV | 1.03E-04 | 1.52E-04 | 9.52E-05 |  |  |
| Cluster V | 8.65E-05 | 1.35E-04 | 7.83E-05 | 7.75E-05 |  |

**Table S2.** Amino acids in the SARS-COV-2 genome exchanged with a frequency of more than 1%.

| gene | protein | position | original amino acid | mutant amino acid | frequency of mutation |
| --- | --- | --- | --- | --- | --- |
| ORF1a | nsp2 | 198 | Val | Ile | 2.02% |
|  |  | 212 | Gly | Asp | 3.09% |
|  |  | 559 | Ile | Val | 3.25% |
|  |  | 585 | Pro | Ser | 3.56% |
|  | nsp3 | 58 | Ala | Thr | 2.87% |
|  | nsp4 | 308 | Phe | Tyr | 1.23% |
|  | nsp5 | 15 | Gly | Ser | 1.02% |
|  |  | 90 | Lys | Arg | 1.17% |
|  | nsp7 | 25 | Ser | Leu | 1.37% |
| ORF1ab | nsp13 | 392 | Arg | Cys | 1.00% |
|  | nsp14 | 320 | Ala | Val | 1.14% |
| ORF3 | ORF3 | 196 | Gly | Val | 1.19% |
| M | M | 3 | Asp | Gly | 1.25% |
|  |  | 175 | Thr | Met | 4.03% |
| ORF8 | ORF8 | 62 | Val | Leu | 1.09% |
| N | N | 193 | Ser | Ile | 1.68% |
|  |  | 197 | Ser | Leu | 1.21% |

**Table S3.** Selection analysis of ORF1ab and S of SARS-CoV-2.

| Gene | protein | position | mutation | SLAC | FUBAR | FEL | MEME |
| --- | --- | --- | --- | --- | --- | --- | --- |
| ORF1a | nsp2 | 85<sup>#</sup> | Thr→Ile | <0.01 | >0.99 | <0.01 | <0.01 |
|  | nsp2 | 339 | Gly→Ser | 0.05 | 0.97 | 0.05 | >0.05 |
|  | nsp3 | 142 | Leu→Phe | >0.05 | 0.96 | 0.04 | 0.05 |
|  | nsp3 | 444 | Asn→Thr | >0.05 | 0.99 | 0.02 | 0.03 |
|  | nsp5 | 90 | Lys→Arg | >0.05 | 0.99 | 0.04 | >0.05 |
|  | nsp6 | 37<sup>#</sup> | Leu→Phe | <0.01 | 0.99 | <0.01 | <0.01 |
| ORF1ab | nsp12 | 323<sup>#</sup> | Pro→Leu | 0.03 | 0.99 | 0.034 | 0.02 |
| S | S1 | 614<sup>#</sup> | Asp→Gly | <0.01 | >0.99 | <0.01 | <0.01 |

“ # ” indicates that the site is also a high frequently exchanged amino acid.
p values for SLAC, FEL and MEME $\leq 0.01$ , FUBAR posterior probability $\geq 0.99$ are considered as putative positive selection sites.

## Figure Legends

**Figure S1: Whole genome phylogenetic tree showing the distribution of the high-frequency mutation sites in each cluster.** The five clusters are color coded, as shown in the inset. The semi-circular arches show the distribution of the high frequency mutation sites in the clusters, each mutation is highlighted with a different colour, as shown in the inset. The bootstrap values at each node of the tree are also shown.

**Figure S2: Phylogenetic tree of S including the location of the high frequency mutation site D614G which revealed some signal for positive selection.** The outer circle represents the origin of a sequence from a certain continent, and the inner circle from a certain country, as specified at the left. The triangle-shaped shadow represents the position of the mutant strains in the phylogenetic tree.

