## Supplementary figures and images for "Snapshot of the evolution and mutation patterns of SARS-CoV-2"

### Fig S1.pdf

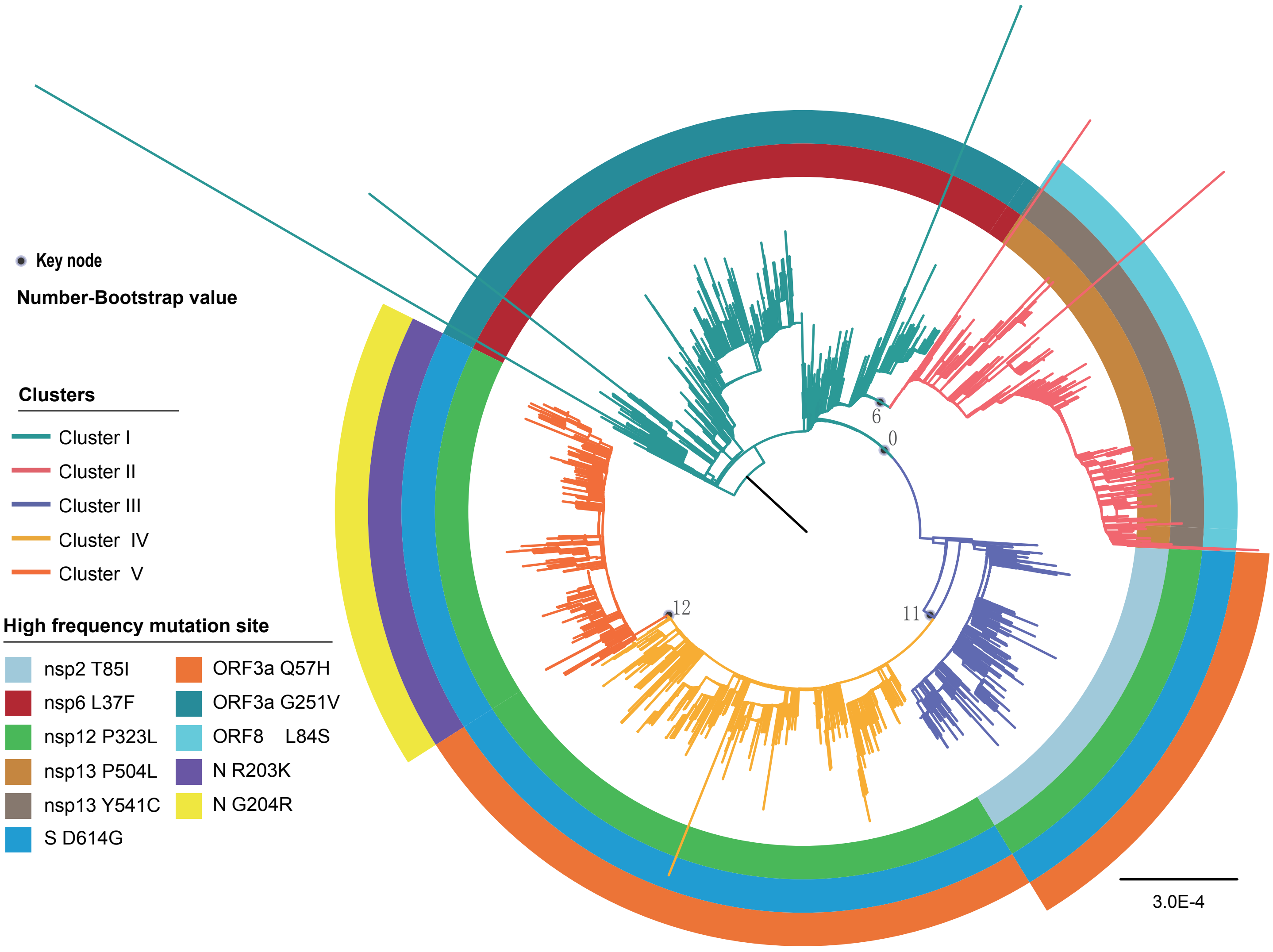

### Fig S2.pdf

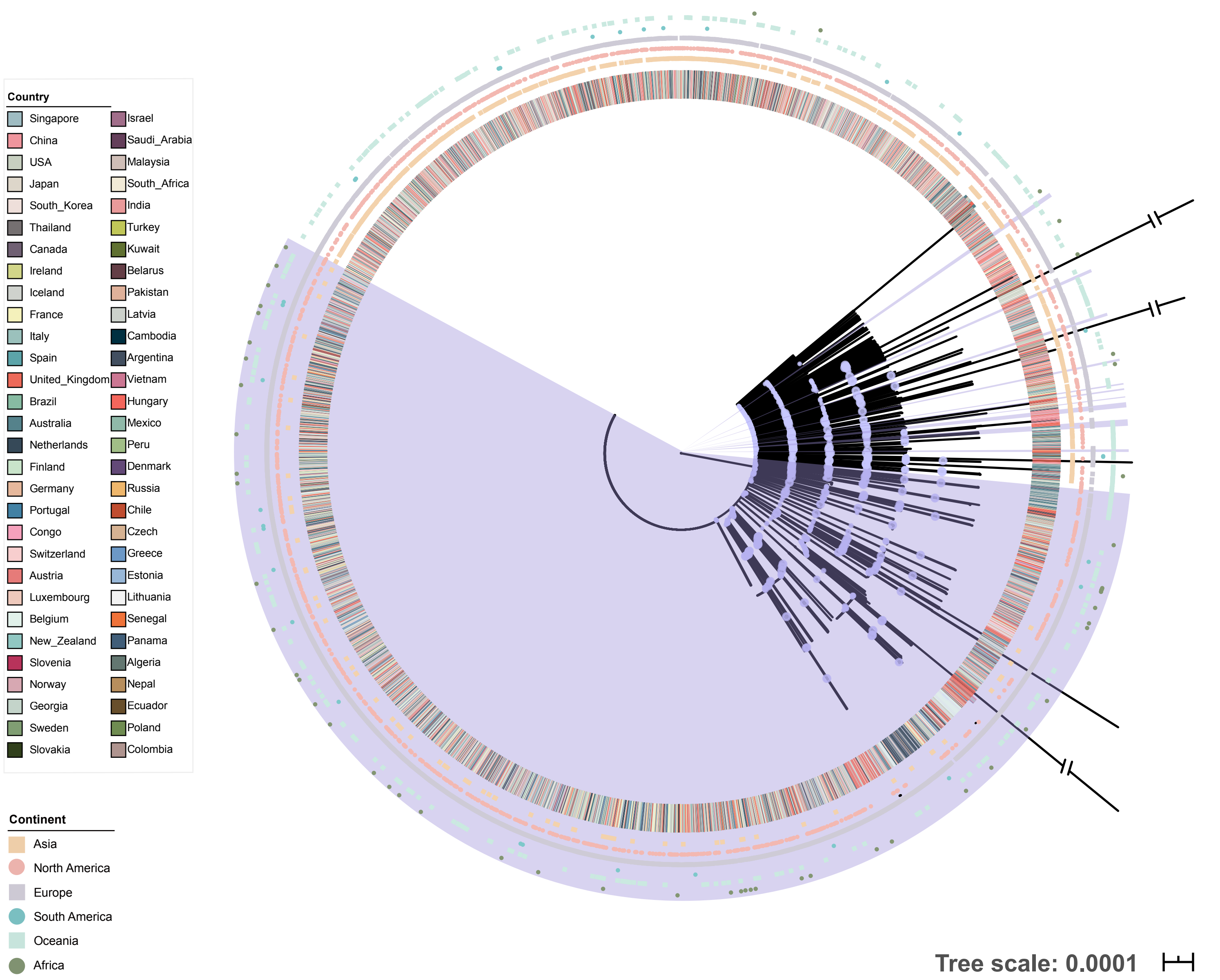
